# The lysosomal V-ATPase B1 subunit in renal proximal tubule is linked to nephropathic cystinosis

**DOI:** 10.1101/2020.07.24.219808

**Authors:** Amer Jamalpoor, Albertien M van Eerde, Marc R Lilien, Charlotte AGH van Gelder, Esther A Zaal, Floris A Valentijn, Roel Broekhuizen, Eva Zielhuis, Julia E Egido, Maarten Altelaar, Celia R Berkers, Rosalinde Masereeuw, Manoe J Janssen

## Abstract

Variants in ATP6V1B1, the gene encoding the B1 subunit of the vacuolar H+-ATPase lead to distal renal tubular acidosis and hearing loss of variable degree. Apart from metabolic acidosis, a 23-month-old girl with pathogenic ATP6V1B1 variants presented with increased granulocyte cystine levels, and symptoms of renal Fanconi syndrome, indicating an unknown link to proximal tubulopathy and cystinosis, a disorder of the proximal tubule caused by variants in the CTNS gene. Here, we demonstrated that ATP6V1B1 is expressed in proximal tubules of human kidney tissue, though to a lesser extent than in distal tubules. Further, we used CRISPR/Cas9 technology to selectively knockout ATP6V1B1 or CTNS in human renal proximal tubule cells and performed a full metabolomic and proteomic analysis to compare their phenotype to isogenic control cells. In line with the clinical data, the loss of ATP6V1B1 disrupted the chemiosmotic coupling between the lysosomal V ATPase B1 subunit and cystinosin and resulted in intralysosomal accumulation of cystine and autophagy activation in renal proximal tubule cells. In conclusion, we identified ATP6V1B1 as a central player in renal proximal tubule cells, regulating lysosomal cystine transport and autophagy, and its absence can lead to proximal tubule dysfunction.

## Introduction

A 23-month-old girl presented with increased granulocyte cystine levels on top of metabolic acidosis and signs of renal Fanconi syndrome. With a working diagnosis of nephropathic cystinosis, treatment with cysteamine, a cystine depleting agent, and electrolytes was started, in parallel to genetic testing. The treatment markedly improved the patient’s clinical conditions and laboratory values; her cystine levels dropped below the reference value, which is considered unusual for a patient with nephropathic cystinosis.

Nephropathic cystinosis (MIM_219800) is an autosomal recessive lysosomal storage disease caused by pathogenic variants in the *CTNS* gene, which encodes the lysosomal H^+^/cystine symporter, cystinosin [1]. The disease is characterized by the accumulation of cystine throughout the body, causing irreversible damage to various organs, particularly the kidneys [2]. Clinically, nephropathic cystinosis patients first develop renal Fanconi syndrome, a general renal proximal tubular dysfunction characterized by phosphaturia, glucosuria, impaired bicarbonate reabsorption and generalized aminoaciduria [2, 3]. With time, this progresses to other complications such as growth retardation, rickets, and chronic kidney disease. The main treatment option currently available is cysteamine, an aminothiol that reduces cystine by converting it into cysteine and cysteine-cysteamine mixed disulfide, both of which are able to exit the lysosomes [4]. Although this drug is found effective in depleting cystine, it is not efficient to revert the signs associated with renal Fanconi syndrome [5].

The diagnostic renal gene panel sequencing revealed pathogenic variants in the *ATP6V1B1* gene (NM_001692.3), while the *CTNS* gene was found to be intact. The *ATP6V1B1* gene codes for the kidney isoform of the B1 subunit of the V1 domain of the vacuolar H^+^-ATPase (V-ATPase; ATP6V1B1), responsible for acidification of intracellular organelles and maintaining renal acid-base homeostasis. Variants in this gene lead to autosomal recessive distal renal tubular acidosis (dRTA) with sensorineural deafness [6, 7]. Moreover, patients with distal renal tubular acidosis and *ATP6V1B1* gene variants have also shown to display proximal tubular dysfunction that resolved after alkali treatment [8]. However, increased cystine levels were not reported, suggesting a potential role for this gene in proximal tubulopathy, as well as a possible link to cystinosis pathophysiology.

Both ATP6V1B1 and cystinosin can be found in the lysosomal membrane of kidney cells and may cooperate in several ways. As a subunit of the V-ATPase, ATP6V1B1 is thought to play a role in lysosome acidification, which in turn is important for the function of cystinosin that depends on this proton gradient for the export of protons and cystine from the lysosome into the cytosol [9, 10]. Besides this chemiosmotic coupling, both cystinosin and the V-ATPase complex closely interact with mammalian target of rapamycin complex 1 (mTORC1), regulating autophagy [9, 11, 12]. Intriguingly, different variants of V-ATPases are also found to regulate transcription factor EB (TFEB), a master regulator of autophagy [13], whose function is also dysregulated in cystinosis [14, 15].

In this study, we aimed to investigate the role of the B1 subunit of the lysosomal V-ATPase in proximal tubules and compare *ATP6V1B1* loss to *CTNS* loss, which is the most common cause of renal Fanconi syndrome in children and has been well studied in proximal tubule cells. We hypothesized that renal proximal tubular acidosis leads to cystinosis-like features. To this end, we confirmed the presence of ATP6V1B1 in human kidney tissue and developed CRISPR-generated *ATP6V1B1*-deficient proximal tubule cells, serving as a disease model, and compared their phenotype to well-characterized isogenic *CTNS^-/-^* cells [14]. Furthermore, we applied a dual-omics approach, combining proteomics and metabolomics, to bridge the gap between the *ATP6V1B1* gene defect and cystinosis by identifying the pathways commonly affected in cystinosis and acidosis proximal tubule cells.

## Results

### Patient with *ATP6V1B1* variant displays a cystinosis-like phenotype of increased cystine levels, metabolic acidosis and symptoms of renal Fanconi syndrome

A 23-month-old girl presented with symptoms of failure to thrive (height Z-score of -2.5 and weight Z-score of -2), and abnormal gait. The patient had impaired hearing from infancy. This was attributed to an enlarged vestibular aqueduct, demonstrated by CAT-scan at the age of four months.

Upon physical examination, she showed plump knee joints with varus deformity. Laboratory examination showed a normal anion-gap metabolic acidosis, hypokalemia, hypophosphatemia, generalized aminoaciduria and elevated cystine levels in granulocytes (Table 1). Renal ultrasound showed bilateral medullary hyper echogenicity and conventional X-ray examination of the knees showed epiphyseal fraying and cupping deformity. Based on the clinical and laboratory findings, the diagnosis of renal rickets due to generalized proximal tubular dysfunction (renal Fanconi syndrome) as a result of nephropathic cystinosis was made. Supplementation with potassium citrate, sodium phosphate, alfacalcidol (active vitamin D) and cysteamine bitartrate (Cystagon^®^) was started. This resulted in an improvement of the patient’s clinical conditions. However, diagnostic sequencing and multiplex ligation-dependent probe amplification did not show variants in the *CTNS* gene. Pending further genetic analysis, treatment with cysteamine bitartrate was continued. Cysteamine bitartrate treatment completely normalized cystine levels in granulocytes (0.03 nmol/mg), which was considered very unusual for a patient with nephropathic cystinosis. Further, ophthalmologic examination did not show corneal deposition of cystine crystals. Cysteamine bitartrate treatment was therefore withheld, after which cystine levels in granulocytes increased slightly to levels below those reported in heterozygous carriers of *CTNS* variants (0.22 nmol/mg). Diagnostic renal gene panel (a panel of renal genes from whole exome sequencing data) analysis was performed, which reiterated the negative result in *CTNS* but demonstrated two heterozygous pathogenic variants in *ATP6V1B1* (NM_001692.3:c.[175-1G>C];[1155dup], p.[(?)];[(Ile386fs)]) [16]. By analyzing the parents, the variants were shown to be in trans.

**Table 1.** Laboratory results of a 23-month-old girl with pathogenic *ATP6V1B1* variants.

| **Parameters** | **Reference** | **Pre-treatment** | **After-treatment** |
| --- | --- | --- | --- |
| **Blood Parameters** |  |  |  |
| pH | 7.35 - 7.45 | 7.28 | 7.36 |
| Bicarbonate (mmol/L) | 22 - 29 | 8.9 | 22.6 |
| Sodium (mmol/L) | 136 - 146 | 141 | 139 |
| Potassium (mmol/L) | 3.8 - 5.0 | 2.9 | 3.9 |
| Chloride (mmol/L) | 99 - 108 | 111 | 105 |
| BUN (mmol/L) | 3.0 - 7.5 | 5.8 | 6.6 |
| Creatinine (μmol/L) | 11 - 34 | 30 | 29 |
| Ionized calcium (mmol/L) | 1.15 - 1.32 | 0.96 | 1.32 |
| Magnesium (mmol/L) | 0.70 - 1.00 |  | 0.98 |
| Inorganic phosphate (mmol/L) | 1.25 - 2.10 | 0.74 | 1.74 |
| Glucose (mmol/L) | 3.6 - 5.6 | 4.8 | 4.4 |
| Cystine (nmol/mg protein) | < 0.17 | 1.02 | 0.22 |
| **Urine Parameters\*** |  |  |  |
| Creatinine (mmol/L) | NA | - | 1.3 |
| Sodium (mmol/L) | NA | - | 49 |
| Potassium (mmol/L) | NA | - | 50 |
| Calcium (mmol/L) | NA | - | < 1.00 |
| Phosphate (mmol/L) | NA | - | 11.7 |
| TmP/GFR (mmol/L) | 1.31 - 1.73 | - | 1.48 |
| Glucose (mmol/L) | NA | - | < 1.00 |
| Citrate (mmol/L) | 0.6 - 4.8 | - | 0.2 |
| Albumin (mg/L) | < 30 | - | 8 |
| Albumin/creatinine ratio (mg/mmol) | < 2.5 | - | 5.9 |
| β2 microglobulin (mg/L) | < 0,2 | - | 94 |
| β2 MG/creatinine ratio (mg/mmol) | < 0.04 | - | 72.3 |
| **Urine Metabolites (mmol/mol creatinine)** |  |  |  |
| Free sialic acid | 16 - 44 | 28 | - |
| Total sialic acid | 52 - 126 | 118 | - |
| Bound sialic acid | 33 - 83 | 90 | - |
| *Taurine* | 12 - 159 | 175 | - |
| *Asparaginic acid* | 3 - 10 | 30 | - |
| *Threonine* | 15 - 62 | 888 | - |
| *Serine* | 45 - 124 | 1045 | - |
| *Asparagine* | < 32 | 475 | - |
| *Glutamic acid* | < 11 | 74 | - |
| *Glutamine* | 62 - 165 | 1895 | - |
| *Proline* | < 13 | < 24 | - |
| *Glycine* | 110 - 356 | 468 | - |
| *Alanine* | 41 - 130 | 157 | - |
| *Citrulline* | < 7 | 298 | - |
| *Alpha-aminobutyric acid* | < 8 | 49 | - |
| *Valine* | 7 - 21 | 312 | - |
| *Cysteine* | 5 - 13 | 166 | - |
| *Isoleucine* | < 6 | 45 | - |
| *Leucine* | 3 - 17 | 124 | - |
| *Tyrosine* | 13 - 48 | 474 | - |
| *phenylalanine* | 10 - 31 | 180 | - |
| *Ornithine* | < 8 | 171 | - |
| *Lysine* | 16 - 69 | 1034 | - |
| *Histidine* | 87 - 287 | 1025 | - |
| *Arginine* | < 8 | 66 | - |

\*, There were no results obtained on urine biochemistry prior to the treatment as it was not possible to collect urine; NA, Not applicable; TmP/GFR, Tubular maximal phosphate reabsorption related to glomerular filtration rate.

With these findings, a definite diagnosis of distal renal tubular acidosis and deafness (OMIM #267300) was established. Sodium phosphate and alfacalcidol supplementation was stopped, while treatment with potassium citrate was continued. The treatment resolved hypophosphatemia, generalized aminoaciduria, rickets, and improved the height Z-score to -0.65. However, the patient still presented a mildly increased urinary excretion of β2-microglobulin (Table 1), suggesting a mild defect in proximal tubules. This prompted us to further investigate the possible link between *ATP6V1B1* gene deficiency and the occurrence of a generalized proximal renal tubular defect.

### V-ATPase B1 subunit is expressed along the human distal and proximal segments of the nephron

To study the role of the V-ATPase B1 subunit in proximal tubules, we first assessed its expression in healthy human kidney tissue by immunohistochemistry. As shown in Figure 1A, ATP6V1B1 is expressed in both proximal and distal tubules although with different levels of intensity. High-intensity staining was observed mainly in distal tubules, whereas low-intensity staining was mostly located to proximal tubules. In contrast, no staining was observed in the glomeruli.

**Figure 1.**
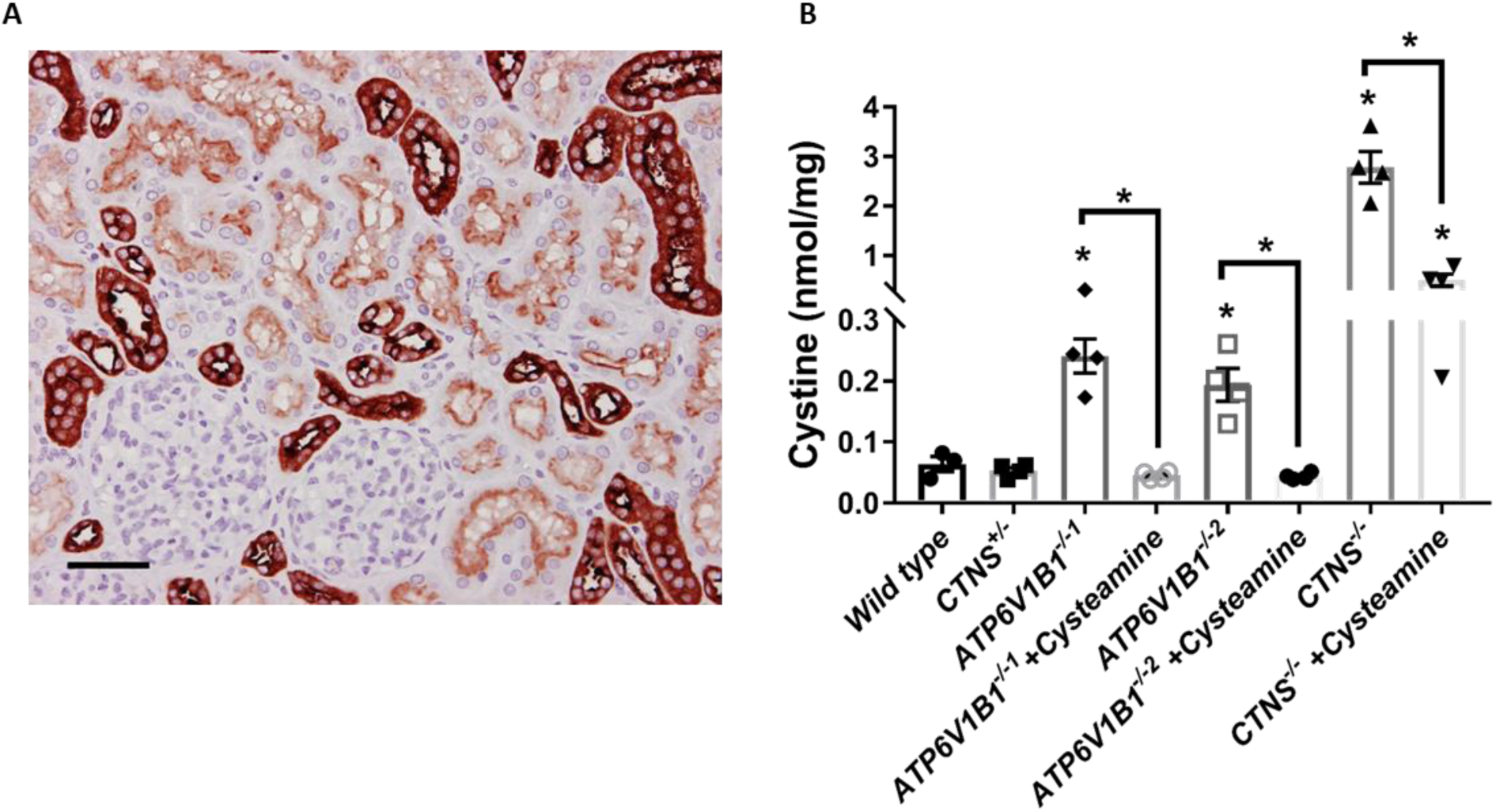
The V-ATPase B1 subunit is expressed in human kidney proximal tubules and regulates cystinosin. **(A)** Immunostaining for the V-ATPase B1 subunit (ATP6V1B1) in human kidney tissue with the ATP6V1B1 antibody. Scale bar is 50 µm. **(B)** Quantification of cystine levels (nmol/mg protein) by HPLC-MS/MS in wild type (control), two clones of *ATP6V1B1^-/-^*, *CTNS^+/-^*, and *CTNS^-/-^*cells treated with or without cysteamine (100 µM). Data are expressed as the mean ± SEM of at least three independent experiments performed in triplicate. P-values < 0.05 were significant.

### CRISPR-generated *ATP6V1B1^-/-^* ciPTEC display increased cystine accumulation and autophagy activation

To dissect the underlying cellular mechanisms of the lysosomal V-ATPase B1 subunit in renal proximal tubules, we introduced a pathogenic variant in the *ATP6V1B1* gene by CRISPR/Cas9 in human renal proximal tubule cells. After cell sorting and subsequent clonal cell expansion, two clones with biallelic variants (*ATP6V1B1^-/-1^*and *ATP6V1B1^-/-2^*) were selected for subsequent experiments. Both *ATP6V1B1*-deficient renal proximal tubule cells displayed a significant increase in cystine levels (∼3-fold), indicating that cystinosin transport function is reduced in absence of the lysosomal V-ATPase B1 subunit (Figure 1B). Nevertheless, the cystine levels were not as high as those found in the isogenic *CTNS^-/-^* cells (45-fold increase compared to control) (Figure 1B). Consistent with the patient data, cysteamine treatment completely normalized cystine levels in *ATP6V1B1^-/-^* cells, but not in *CTNS^-/-^* cells (Figure 1B). Cystine levels in *CTNS^-/-^*cells treated with cysteamine was still higher (7.5-fold) than that found in control cells (Figure 1B).

When looking at the mRNA levels, we found a small (less than 2-fold) reduction in *CTNS* mRNA expression in the *ATP6V1B1* deficient cells (Supplementary figure S1A). It is not expected that this change in mRNA will directly affect the cystine transport function, especially as the same reduction in mRNA is seen in the proximal tubule cells bearing a heterozygous *CTNS* mutation (*CTNS^+/-^* cells), which does not result in cystine accumulation (Supplementary figure S1A, figure 1B). This indicates that the accumulation of cystine in *ATP6V1B1^-/-^* cells is not due to a lack in cystinosin protein levels, but rather caused by a reduced efficiency in the cystinosin activity. It is worth mentioning that *CTNS^-/-^* cells also show a decreased *ATP6V1B1* mRNA expression (1.5-fold) compared to control cells (supplementary figure S1B), suggesting there may be a regulatory link between the expression of these genes.

The lysosomal V-ATPase is a component of the mTOR complex and is involved in autophagy regulation [9]. Disruption of the mTORC1 complex and its dissociation from lysosomes is correlated with TFEB nuclear translocation and induction of autophagy, as also seen in cystinosis [14]. In agreement with reduced mTOR activity, a ∼3.0-fold increase in TFEB nuclear translocation was observed in both *ATP6V1B1* knockout lines compared to control cells, comparable to *CTNS^-/-^* cells (2.3-fold; Figure 2A, 2B). In agreement with TFEB known to downregulate its own expression after activation, we found that the endogenous *TFEB* mRNA expression was reduced in all three lines (∼1.5-fold) when compared to control cells (Figure 2C). During autophagy, LC3-II (Microtubule-associated protein 1A/1B-light chain 3) is recruited to autophagosomes and its accumulation is correlated with abnormal induction of autophagy.[17] In accordance, we found significantly increased LC3-II levels in both the *CTNS* and *ATP6V1B1* knockout cells compared to control cells (Figure 2D). We also assessed the effect of the *ATP6V1B1* gene defect on endocytosis and lysosomal-cargo degradation in the knockout lines using BSA and DQ-BSA uptake, respectively. *ATP6V1B1* knockout lines maintained endocytosis and displayed increased endocytic cargo processing function (Figure 2E, 2F). *CTNS^-/-^* cells, similar to *ATP6V1B1^-/-^* cells, maintained endocytosis functionality but, in contrast displayed a reduced ability (∼2.5-fold) to degrade endocytic cargo compared to control cells (Figure 2E, 2F).

**Figure 2.**
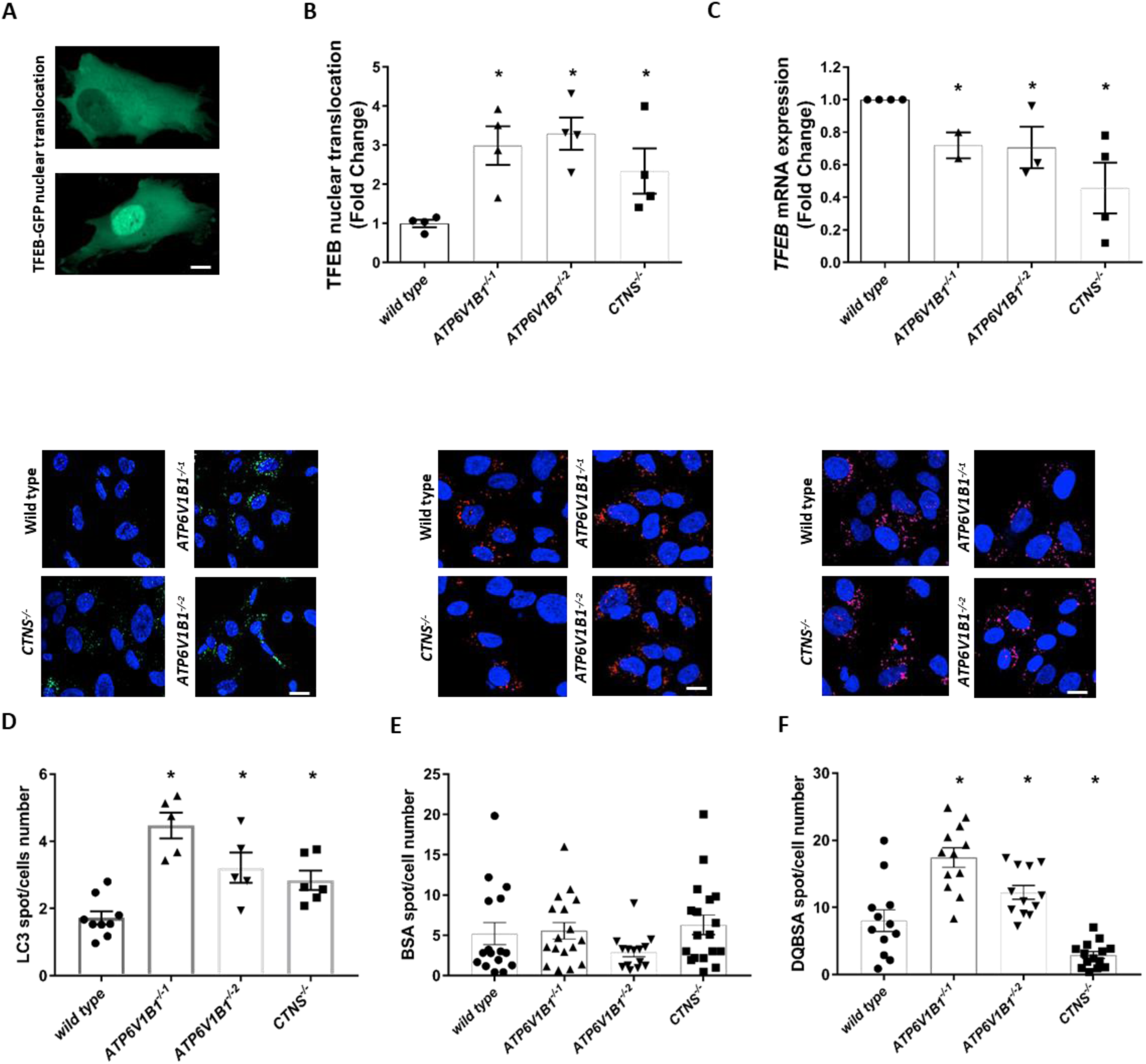
The V-ATPase B1 subunit regulates autophagy without influencing the lysosomal vesicular trafficking in human renal proximal tubule cells. **(A)** Representative confocal micrographs of transcription factor EB (TFEB) cellular distribution. Scale bars is 20 µm. **(B)** Quantification of TFEB nuclear translocation in wild type (control), two clones of *ATP6V1B1^-/-^*, and *CTNS^-/-^* cells, respectively. **(C)** The levels of *TFEB* mRNA expression in control, two clones of *ATP6V1B1^-/-^*, and *CTNS^-/-^* cells. **(D)** Representative confocal micrographs and quantification of LC3-II accumulation in control, two clones of *ATP6V1B1^-/-^*, and *CTNS^-/-^* cells in presence of 25 nM bafilomycin (BafA1) for 4 hrs. Scale bars are 20 µm. **(E, F)** Representative confocal micrographs and quantification of BSA and DQ-BSA in control, two clones of *ATP6V1B1^-/-^*, and *CTNS^-/-^* cells, respectively. Scale bars are 20 µm. Data are expressed as the mean ± SEM of at least three independent experiments performed in triplicate. P-values < 0.05 were significant.

Together, our data indicate that in renal proximal tubule cells, ATP6V1B1 plays a role in autophagy activation and lysosomal cystine transport, without having an effect on the lysosomal vesicular trafficking and degradation.

### Metabolic and proteomic profiling link the lysosomal V-ATPase B1 subunit to nephropathic cystinosis

To further investigate the role of the B1 subunit of the lysosomal V-ATPase in proximal tubules and its contribution to cystinosis pathophysiology, we performed semi-targeted metabolomics and untargeted proteomics in *ATP6V1B1^-/-^*, *CTNS^-/-^* and control cells (Figure 3, 4). Principal component analysis (PCA) of the 100 measured metabolites and over 3,474 identified proteins (Figure 3A, 4A) demonstrated that both *CTNS^-/-^* and *ATP6V1B1^-/-^* cells share quite similar metabolic and proteomic profiles and are different from the control cells. This was further visualized by unsupervised hierarchical clustering in which *ATP6V1B1^-/-^* cells clustered with *CTNS^-/-^* cells but not with control cells (Figure 3B). Moreover, pathway enrichment analysis of the metabolites revealed multiple similarly affected pathways in the two diseased cells, including the arginine and proline metabolism, the ß-alanine metabolism, and the alanine, aspartate and glutamate metabolism (Figure 3C, 3D), providing additional evidence that defects in both genes result in similar metabolic disruptions in proximal tubule cells.

**Figure 3.**
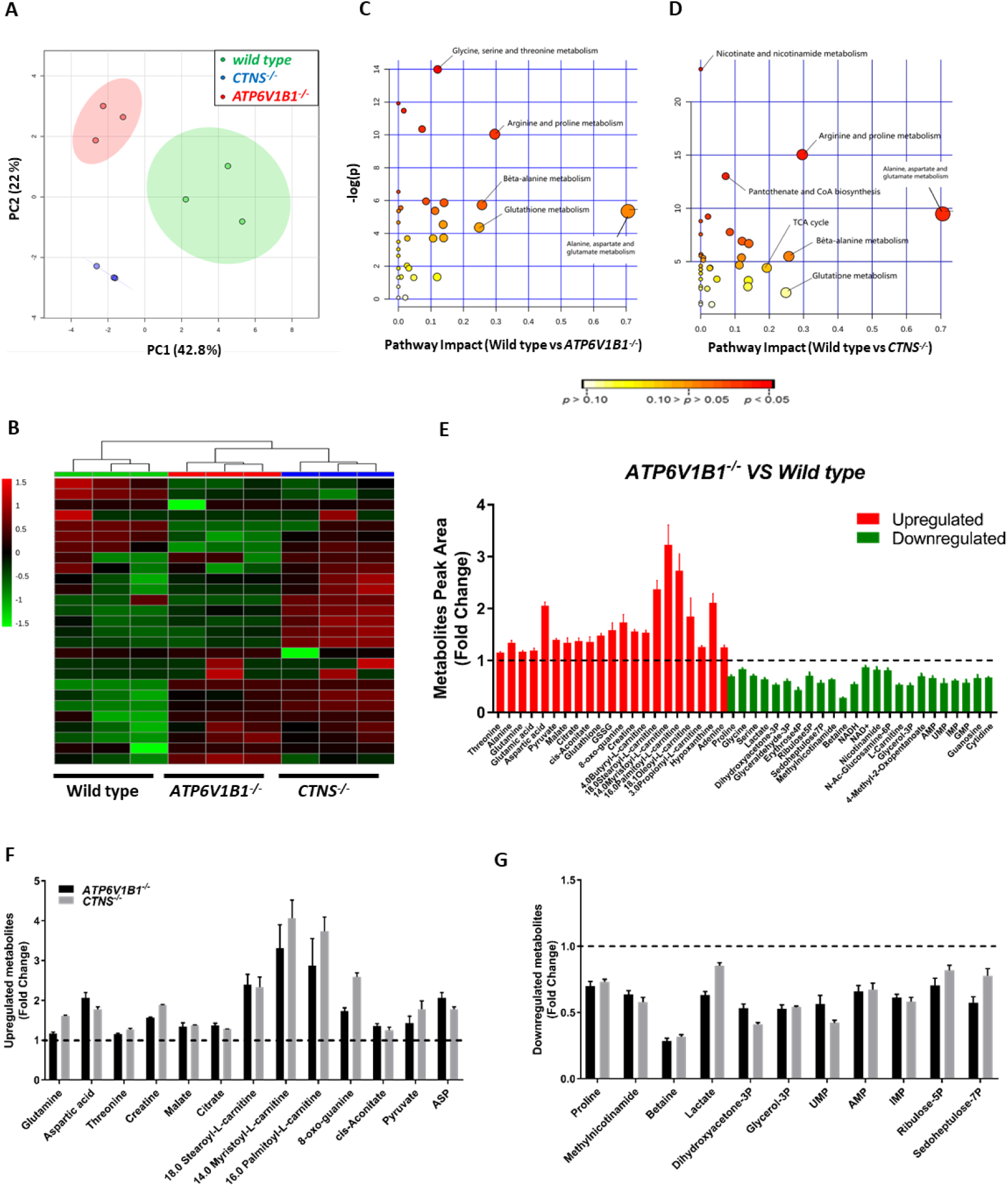
Defects in both *ATP6V1B1* and *CTNS* genes result in similar metabolic disruptions in human renal proximal tubule cells. **(A)** Principal component analysis (PCA) of wild type (control), *ATP6V1B1^-/-^*, and *CTNS^-/-^* cells based on the 100 metabolites measured. **(B)** Heatmap analysis of metabolites distinctively expressed in control, *ATP6V1B1^-/-^*, and *CTNS^-/-^*cells. Color code: green lower than control, red higher than control (p<0.05). **(C, D)** Global test pathway enrichment analysis of the intracellular metabolic interactions distinctively affected in *ATP6V1B1^-/-^* and *CTNS^-/-^* cells compared to control cells, respectively. Larger circles further from the y-axis and orange-red color show higher impact of pathway affected in cells. **(E)** List of metabolites that were significantly altered in *ATP6V1B1^-/-^* compared to control cells (P<0.05). **(F, G)** List of metabolites that were shared and significantly upregulated and downregulated in both *ATP6V1B1^-/-^* and *CTNS^-/-^* cells compared to control cells, respectively. Data are expressed as the mean ± SEM of one experiment performed in triplicate. P-values < 0.05 were significant.

**Figure 4.**
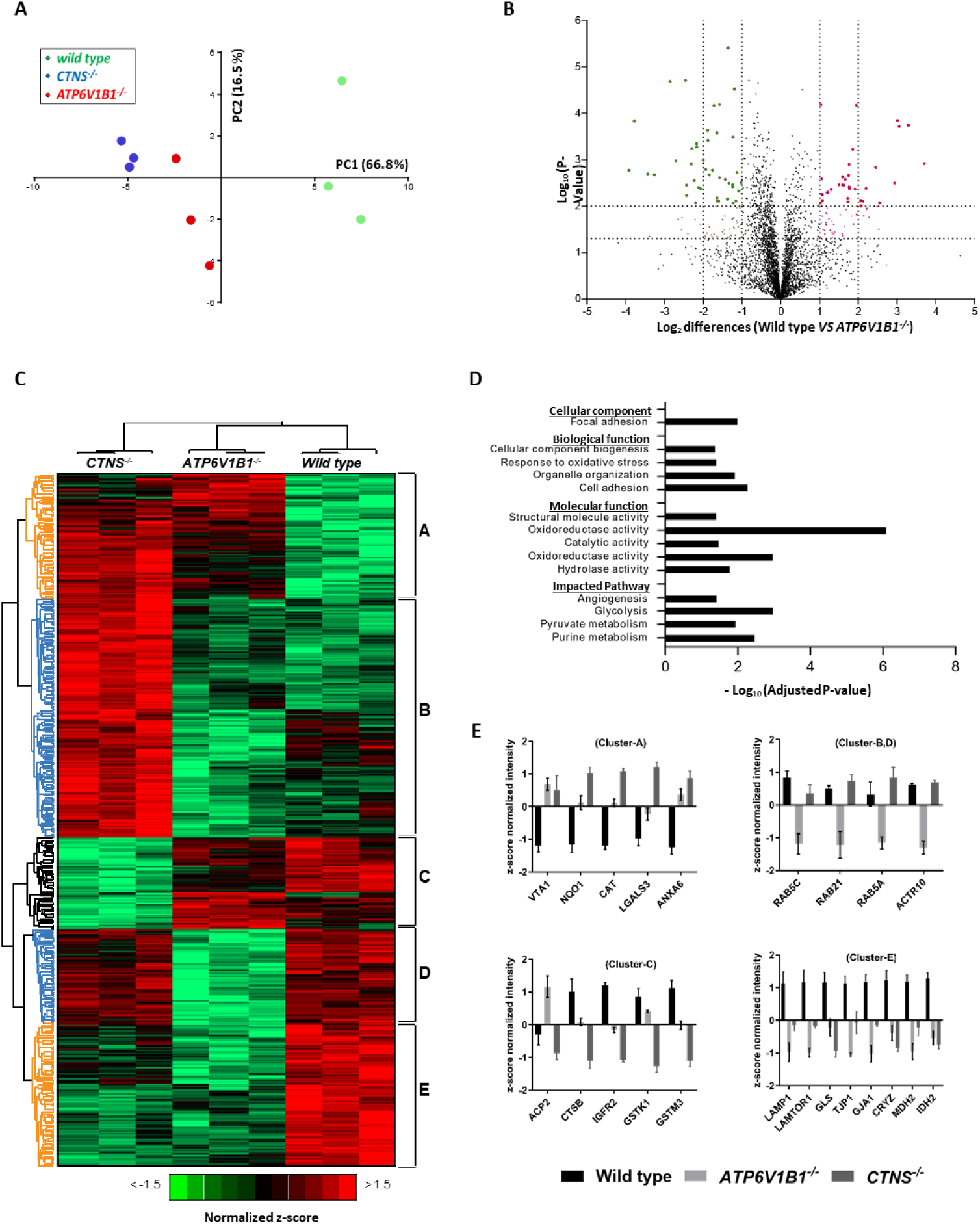
Proteomic profiling bridges the lysosomal V-ATPase B1 subunit to nephropathic cystinosis. **(A)** Principal component analysis (PCA) of the measured proteins in wild type (control), *ATP6V1B1^-/-^*, and *CTNS^-/-^* cells. **(B)** Volcano plot illustrates significantly differentially abundant proteins. The -log10 (Benjamini–Hochberg corrected P-value) is plotted against the log2 (fold change: wild type/ *ATP6V1B1^-/-^*). The non-axial vertical lines denote ±1.5-fold change while the non-axial horizontal line denotes P=0.05, which is our significance threshold (prior to logarithmic transformation). **(C)** Heatmap analysis of the proteins distinctively expressed in control, *ATP6V1B1^-/-^*, and *CTNS^-/-^* cells. The row displays protein feature and the column represents the samples. The row Z-score of each feature is plotted in red-green colour scale. Proteins significantly decreased were displayed in green, while metabolites significantly increased were displayed in red. Hierarchical clustering of these proteins resulted in five main protein clusters. Clusters A and E (in orange) represent proteins that were similarly affected in both *ATP6V1B1^-/-^* and *CTNS^-/-^* cells compared to control cells, clusters B and D (in blue) show proteins affected differently in the *ATP6V1B1^-/-^* as compared to the *CTNS^-/-^* and control cells, and cluster C (in black) indicates proteins affected differently in the *CTNS^-/-^* as compared to *ATP6V1B1^-/-^* and control cells. **(D)** Gene ontology analysis of the significantly changed proteins in cluster-A, E and cluster-B, D of the Heatmap. **(E)** List of proteins that were significantly altered in *ATP6V1B1^-/-^*, and *CTNS^-/-^* cells compared to control cells. Data are expressed as the mean ± SEM of one experiment performed in triplicate. P-values < 0.05 were significant.

To unravel the role of the lysosomal V-ATPase B1 subunit in proximal tubules, we first evaluated metabolite and protein expression differences between *ATP6V1B1^-/-^* and control cells (Figure 3E, 4B), and the differentially expressed metabolites and proteins were then compared with those in *CTNS^-/-^* cells. About 50% of the quantified metabolites (especially amino acids that are involved in acid regulation) [18, 19] were significantly altered as a result of ATP6V1B1 loss in proximal tubule cells (Figure 3E). Cross-checking individual metabolites that were similarly affected in both *ATP6V1B1^-/-^* and *CTNS^-/-^*cells, revealed that *CTNS^-/-^* cells also present a fairly similar metabolite alteration as those found in *ATP6V1B1^-/-^* cells (Figure 3F, 3G). Proteomic profiling, on the other hand, revealed a total of 562 proteins that were differentially expressed in *ATP6V1B1^-/-^* and *CTNS^-/-^* cells compared to controls (Figure 4C). Hierarchical clustering of these proteins resulted in five main protein clusters. Clusters A and E (in orange) represent proteins that were similarly affected in both *ATP6V1B1^-/-^* and *CTNS^-/-^* cells compared to control cells, clusters B and D (in blue) show proteins affected differently in the *ATP6V1B1^-/-^* as compared to the *CTNS^-/-^* and control cells, and cluster C (in black) indicates proteins affected differently in the *CTNS^-/-^* as compared to *ATP6V1B1^-/-^*and control cells. The differentially abundant proteins were then subjected to gene ontology classification via the Panther Classification System database [20] to highlight their biological and molecular functions, and cellular component in the cells (Figure 4D). The analysis showed an overall reduction in proteins involved in catalytic activity, oxidoreductase activity, cell-cell adhesion, organelle organization, and focal adhesion in *ATP6V1B1^-/-^* and *CTNS^-/-^* cells compared to controls. Key proteins involved in lysosomal cargo degradation, namely lysosomal acid phosphatase (ACP2), Cathepsin B (CTSB), and mannose 6-phosphate receptors (M6PRs; IGFR2, receptors responsible for the delivery of newly synthesised lysosomal enzymes from Golgi to the lysosome), that were shown to be dysregulated in *CTNS^-/-^* cells [14], were not affected in *ATP6V1B1*^-/-^ cells (Figure 4E-Cluster-C). However, lysosome-associated membrane glycoprotein-1 (LAMP1), and Ragulator complex protein LAMTOR1 were found significantly downregulated in both *ATP6V1B1^-/-^* and *CTNS^-/-^* cells compared to controls (Figure 4E-Cluster-E). *ATP6V1B1^-/-^* cells also showed a decreased expression of several autophagy-regulatory proteins, Rab proteins (Figure 4E-Cluster-B, D), confirming that the loss of ATP6V1B1 in proximal tubule cells results in abnormal autophagy activation, but without influencing the lysosomal cargo degradation. *ATP6V1B1^-/-^* and *CTNS^-/-^* cells also presented decreased expression of tight junction protein, Zonula occludin-1 (TJP1; ZO-1) and gap junction alpha-1 protein (GJA1), and increased expression of lectin and β-galactoside-binding protein family 21, galectin-3 (LGALS3; Figure 4E-Cluster-A), markers of kidney disease progression. Altogether, our findings revealed a previously unrecognized role of the lysosomal V-ATPase B1 subunit in renal proximal tubular epithelial cells and signified its link to cystinosis pathophysiology.

## Discussion

In this work, we revealed a previously unrecognized role of the lysosomal V-ATPase B1 subunit in renal proximal tubular epithelial cells and showed its loss could have similar effects on the cells as *CTNS* loss. In line with the clinical suggestion, the *ATP6V1B1* gene defect in renal proximal tubule cells hampered cystinosin regulation and resulted in cystine accumulation, which was completely normalized by cysteamine. The loss of ATP6V1B1 also entailed drastic cellular changes with a marked induction of autophagy and metabolic acidosis in proximal tubule cells, without influencing the lysosomal vesicular trafficking.

V-ATPases are large multisubunit complexes that utilize the energy derived from the hydrolysis of cytosolic ATP to transport protons across biological membranes and maintain the acidic pH of endocytic and secretory organelles [21, 22]. They are ubiquitously distributed on intracellular tubulo-vesicular membranes, and at the plasma membrane in specialized cell types. The B1 subunit of the V-ATPase has been found expressed only in intercalated cells of the kidney [23–26], inner ear [7], ocular ciliary epithelium [27], male reproductive tract [28], and placenta [29]. The functional importance of the *ATP6V1B1* gene in kidney and auditory physiology is evidenced by the fact that mice and humans with variants in this gene develop the autosomal recessive dRTA with deafness [6, 7, 30]. However, the fact that our patient with *ATP6V1B1* variants also developed symptoms of cystinosis and renal Fanconi syndrome, demonstrated by increased granulocyte cystine levels, generalized aminoaciduria, and urinary excretion of β2-microglobulin, prompted us to investigate the possible link between *ATP6V1B1* gene deficiency and the occurrence of a generalized renal proximal tubular defect.

We demonstrated that ATP6V1B1 is expressed in proximal tubules of human kidney tissue, though to a lesser extent than in distal tubules. In addition, when we specifically knocked out the *ATP6V1B1* gene in human renal proximal tubule cells, we found increased cystine levels with autophagy activation. Cystinosin is a H^+^/cystine symporter that requires lysosomal H^+^ influx mediated by the lysosomal V-ATPase to transport cystine to the cytoplasm [10]. The loss of ATP6V1B1 likely disrupted the chemiosmotic coupling between the lysosomal V-ATPase B1 subunit and cystinosin and resulted in intralysosomal accumulation of cystine in proximal tubule cells (Figure 5). Using an omics-based strategy, we provided evidence that defects in both genes may cause proximal renal tubular acidosis. The proximal tubule is the primary site for active solute reabsorption and secretion, and plays a central role in acid-base balance [31, 32]. During metabolic acidosis, the proximal tubule alters its metabolism and transport properties, leading to increased glutamine levels and renal ammoniagenesis to excrete more acid into urine [18, 33, 34]. In accordance, we observed an accumulation of glutamine in the both diseased cells. This could be due to either increased uptake or decreased catabolism of glutamine as a result of decreased expression of glutamines enzyme (GLS) in *ATP6V1B1* and *CTNS* deficient cells.

**Figure 5.**
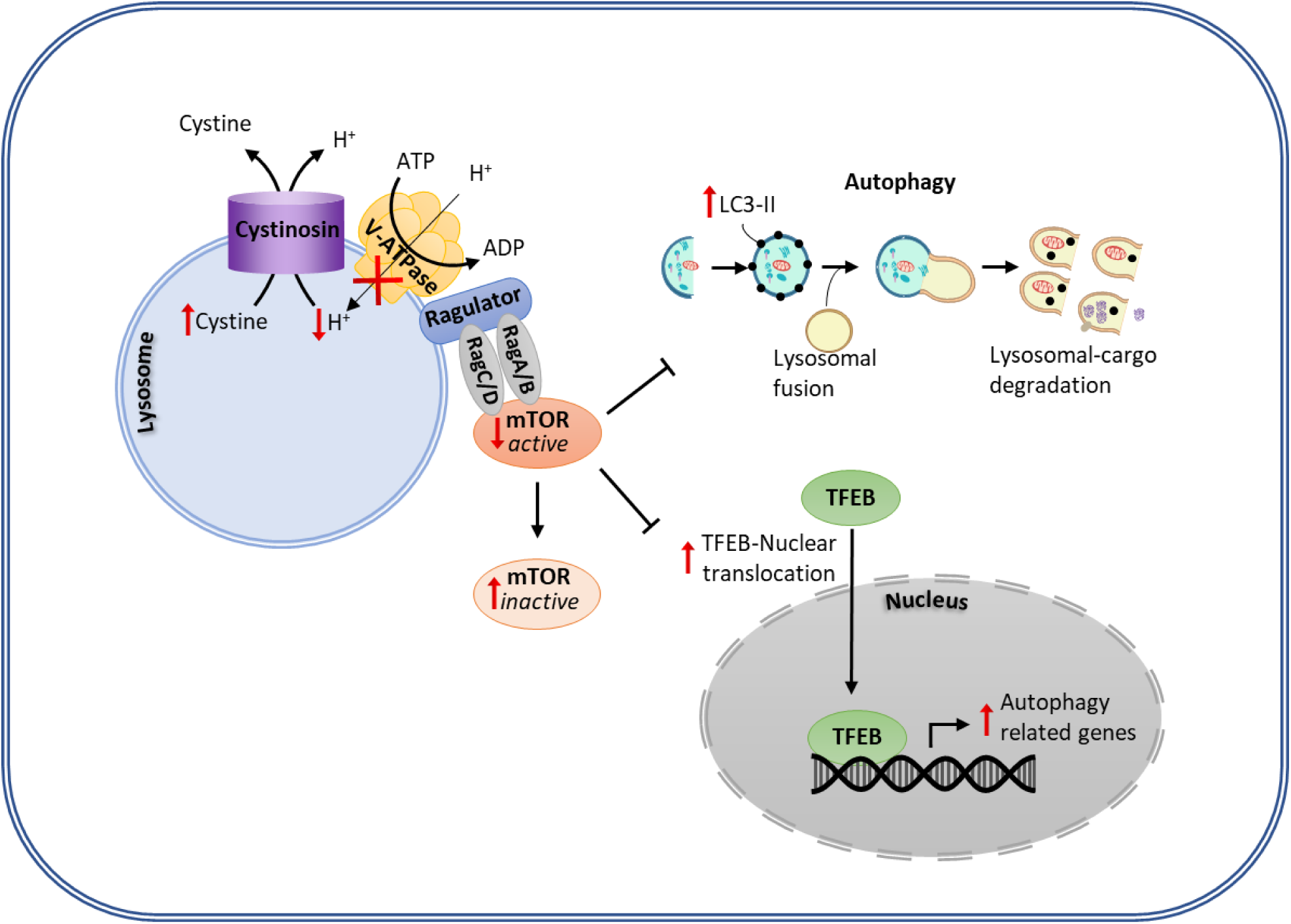
The lysosomal V-ATPase B1 subunit is the central interplay between cystinosin and mTOR-signaling, regulating cystine transport and autophagy in human renal proximal tubule cells. ATP; adenosine triphosphate, ADP; adenosine diphosphate, GTP; Guanosine triphosphate, GDP; Guanosine diphosphate, mTOR; mammalian target of rapamycin complex, LC3-II; Microtubule-associated protein 1A/1B-light chain 3, TFEB; transcription factor EB.

Furthermore, *ATP6V1B1^-/-^* and *CTNS^-/-^* proximal tubule cells showed decreased levels of lactate and increased levels of pyruvate and tricarboxylic acid (TCA) cycle intermediates, namely cis-aconitate, malate and citrate. This is in agreement with the findings of LaMonte *et al.*, where they demonstrated that metabolic acidosis induces aerobic glycolysis, redirecting glucose away from lactate production and towards the TCA cycle in human cancer cells [19]. Proteomic analysis, on the other hand, revealed that the loss of either ATP6V1B1 or cystinosin is accompanied with decreased expression of tight junction protein (ZO-1 and GJA1), indicating proximal tubular epithelial dysfunction and disrupted tight junction integrity.[12, 35, 36] *ATP6V1B1^-/-^* and *CTNS^-/-^* cells also presented increased expression of galectin-3, a protein demonstrated to be markedly upregulated in acute tubular injury [37] and in progressive renal fibrosis [38]. Lobry *et al*., demonstrated that cystinosin loss-of-function results in galectin-3 overexpression in cystinotic mice kidneys, inducing macrophage infiltration and kidney disease progression [39]. In light of our present data, we suggest that the V-ATPase B1 subunit plays an important role in acid-base homeostasis and is involved in the occurrence of a generalized proximal renal tubular defect that could functionally overlap with the one seen in cystinosis.

Nephropathic cystinosis is the most common genetic cause of renal Fanconi syndrome in children. Variants in *CTNS* lead to lysosomal accumulation of cystine, reduced mTOR activation, delayed protein degradation and increased oxidative stress in proximal tubule cells [12, 14, 40–47]. It is known that cystinosin can directly interact with V-ATPase-Ragulator-Rag Complex at the lysosomal membrane, regulating mTOR-mediated autophagy [9]. On the other hand, V-ATPase was also identified as a potential amino acid sensing protein that can recruit and activate mTORC1 [48]. Here, we demonstrated that the defect or the decreased *ATPV1B1* gene expression is corelated with the increased TFEB nuclear translocation and subsequent autophagy activation (Figure 5). Hence, the fact that *CTNS^-/-^* cells have increased autophagy activation could be a direct result of the decreased activity of ATP6V1B1, disabling mTOR recruitment to the lysosomes and subsequent deactivation of mTOR signaling, in line with previous reports demonstrating reduced mTOR activity in *in vitro* and *in vivo* models of cystinosis [9, 12, 14, 49, 50]. This was further evidenced by Ragulator complex protein LAMTOR1 and LAMP1 being downregulated in both *ATP6V1B1^-/-^* and *CTNS^-/-^* cells. LAMTOR1 as a part of V-ATPase-Ragulator-Rag Complex is required for lysosomal recruitment of mTORC1 and consequent activation. Depletion in LAMTOR1 prevents mTOR shuttling to lysosomal surface, resulting in autophagy activation [51, 52]. Taken together, our study identified ATP6V1B1 as a central player in human renal proximal tubule cells, regulating cystine transport and autophagy, and its absence can lead to proximal tubule dysfunction.

## Methods

### Reagents

All chemicals and reagents were obtained from Sigma-Aldrich (Zwijndrecht, The Netherlands) unless specified otherwise. Primary antibodies used were rabbit anti-ATP6V1B1 (Abcam #ab192612, dilution 1:500) and mouse anti-LC3 (Novus Biologicals #NB600-1384SS, dilution 1:1000). Secondary antibodies used were goat anti-rabbit BrightVision Horse Radish Peroxidase (Klinipath, Duiven, The Netherlands) and goat anti-mouse (#A32723, dilution 1:5000) obtained from Dako products (CA, USA).

### Immunohistochemistry on human renal tissue sections

A tumor-free human kidney tissue section was derived from a patient undergoing nephrectomy for renal carcinoma at the University Medical Center Utrecht (UMCU). The patient sample was anonymized. Because leftover material from routine clinical procedures in the UMCU was used, informed patient consent and additional ethical approval were not required. The UMCU policy allows anonymous use of redundant tissue for research purposes as part of the standard treatment agreement with patients in the UMCU [53]. Renal tissue was fixed in a 4% buffered formalin solution for 24 hrs and subsequently embedded in paraffin blocks. Sections of 3 μm were cut and mounted on adhesive slides (Leica Xtra) and rehydrated through a series of xylene and alcohol washes after which slides were rinsed in distilled water. First, endogenous peroxidase was blocked using H_2_O_2_. This was followed by heat-based antigen retrieval in citrate buffer (pH 6) and primary antibody incubation (anti-ATP6V1B1, Abcam ab192612, 1:500) diluted in PBS/1%BSA. After incubation with goat anti-rabbit BrightVision Horse Radish Peroxidase linked secondary antibody (Klinipath, Duiven, The Netherlands), sections were stained using Nova Red substrate (Vector Laboratories, Burlingame, CA, USA) and counterstained with Mayer’s haematoxylin. Images were acquired using a Nikon Eclipse E800 microscope.

### CiPTEC culture

The conditionally immortalized proximal tubular epithelial cells (ciPTEC) (MTA #A16-0147) were obtained from Cell4Pharma (Nijmegen, The Netherlands) and were cultured as described previously by Wilmer et al. 2010 [54]. In short, these cells were immortalized using the temperature sensitive SV40ts A58 and human telomerase gene (hTERT), which allows them to be expanded at 33°C and regain a mature phenotype after culturing at 37°C. The culture medium was Dulbecco’s modified Eagle medium DMEM/F-12 (GIBCO) supplemented with fetal calf serum 10% (v/v), insulin 5 µg/ml, transferrin 5 µg/ml, selenium 5 µg/ml, hydrocortisone 35 ng/ml, epidermal growth factor 10 ng/ml and tri-iodothyronine 40 pg/ml. Cells were seeded at a density of 55,000 cells/cm^2^ and grown at 33°C for 24 hrs to enable them to proliferate and subsequently cultured at 37°C for 7 days to mature into fully differentiated tubular epithelial cells.

### Generation of *ATP6V1B1^-/-^* and *CTNS*^-/-^ isogenic ciPTEC lines

We previously created an isogenic CRISPR-mediated cystinotic ciPTEC line [14] referred to as *CTNS^-/-^*. Using the same parent ciPTEC line, we created the *ATP6V1B1*^-/-^ cell line as outlined. Guide RNAs (gRNAs) targeting exon 4 of the *ATP6V1B1* gene were designed using the online gRNA designing tool available at chopchop.cbu.uib.no. To maximize specificity, guide sequences with high scores for on-target efficiency and no predicted off-targets having at least 3 base pair mismatches in the genome were selected. Optimal gRNA (5’-CCTACGAACTCCGGTGTCAG-3’) was cloned into the pSPCas9(BB)-2A-GFP plasmid (Addgene #48138) as described previously by Ran et al. [55], and introduced into ciPTEC using PolyPlus JetPrime. At 72 hrs post-transfection, GFP-positive singlet cells were sorted using FACS Aria-II flow cytometer and expanded in 96-wells plate. The gRNA cut site was amplified with PCR using the primers flanking the cut region (F. *ATP6V1B1*_ex4 5’-ACTCTGAAGGCAGGAAATGGTC-3; R.*ATP6V1B1*_ex4 5’-GAGGAAGGTGGGTTCAATAAC-3’). Finally, knockouts were confirmed by Sanger sequencing.

### Intracellular cystine quantification by HPLC-MS/MS

Cystine levels were quantified using high-performance liquid chromatography-tandem mass spectrometry (HPLC-MS/MS) using a rapid and sensitive assay developed and validated in house [56]. In brief, cell pellets were suspended in N-Ethylmaleimide (NEM) solution containing 5 mM NEM in 0.1 mM sodium phosphate buffer (pH 7.4). The cell suspension was then precipitated, and protein was extracted with sulfosalicylic acid 15% (w/v) and centrifuged at 20,000 g for 10 min at 4°C. Protein concentration was determined by the method of the Pierce™ BCA protein assay kit according to the manufacturer’s protocol (Thermo Fisher, The Netherlands), and the cystine concentration was measured using HPLC-MS/MS. Data are expressed as the cystine values (nmol) corrected for protein content (mg).

### Quantitative real-time PCR

The mRNAs were extracted from cells using the Qiagen RNeasy mini kit according to the manufacturer’s instructions. Total mRNA (600 ng) was reverse transcribed using iScript Reverse Transcriptase Supermix (Bio-Rad). Quantitative real-time PCR was performed using iQ Universal SYBR Green Supermix (Bio-Rad) with the specific sense and anti-sense primers for *ATP6V1B1* (forward: 5’-TGGATATCAATGGCCAGCCC-3’; reverse: 5’-CTTCGCCATCGTCTTTGCAG-3’), *CTNS* (forward: 5’-AGCTCCCCGATGAAGTTGTG-3’; reverse: 5’-GTCAGGTTCAGAGCCACGAA-3’), and *TFEB* (forward: 5’-GCAGTCCTACCTGGAGAATC-3’; reverse: 5’- TGGGCAGCAAACTTGTTCC-3’). The ribosomal protein S13 (RPS-13) (forward: 5’-GCTCTCCTTTCGTTGCCTGA-3’; reverse: 5’- ACTTCAACCAAGTGGGGACG-3’) was used as the reference gene for normalization and relative expression levels were calculated as fold change using the *2^-ΔΔ^Ct* method.

### Immunofluorescence and confocal microscopy

To investigate LC3-II accumulation, cells were seeded in a special optic 96-well plate in presence of 25 nM bafilomycin (BafA1) for 4 hrs. Thereafter, cells were fixed with 4% paraformaldehyde in phosphate buffered saline (PBS) for 10 min, permeabilized with 0.1% Triton-X solution for 10 min, and blocked with 1% bovine serum albumin (BSA) diluted in PBS for 30 min. Subsequently, cells were stained with the primary antibody (mouse anti-LC3, dilution 1:1000) diluted in blocking buffer overnight at 4°C. After 3 washes with PBS, the cells were incubated for 2 hrs at room temperature with the secondary antibody (goat anti-mouse, dilution 1:5000). Nuclei were stained with Hoechst 33342 (1 µM) and cells were imaged using a Cell Voyager 7000 (CV7000) confocal microscope (Yokogawa Electric corporation, Tokyo, Japan).

To assess TFEB intracellular distribution, cells were seeded in a special optic 96-well plate until reaching 50% confluence. Cells were then transfected with the TFEB-GFP plasmid (a kind gift from Dr. Annelies Michiels (Viral Vector Core, Leuven, Belgium)) using PolyPlus JetPrime reagent according to the manufacturer’s instructions. After 48 hrs from transient transfection, cells were stained with Hoechst 33342 (1 µM) for 10 min and imaged using a Cell Voyager 7000 (CV7000) confocal microscope (Yokogawa Electric corporation, Tokyo, Japan). TFEB nuclear translocation data are expressed as number of cells with nucleus-TFEB positive over the total number of TFEB-transfected cells.

### Endocytosis assay

The endocytic uptake was monitored in ciPTEC following incubation for 1.5 hr at 37°C with 50 μg/ml of either BSA-AlexaFluor-647 (A34785, Thermo Fisher Scientific) or DQ Red BSA (D12051, Invitrogen). The cells were then fixed and stained with Hoechst 33342 (1 µM) for 10 min and imaged using a CV7000 confocal microscope (Yokogawa Electric corporation, Tokyo, Japan). Data were quantified with Columbus™ Image Data Storage and analysis software (PerkinElmer, Groningen, The Netherlands). Data are expressed as the number of BSA/DQ Red BSA spots per cell.

### Metabolomics and Proteomics

The omics analyses were performed using high performance liquid chromatography mass spectrometry (HPLC/MS) and the data were analyzed as described previously [14]. The mass spectrometry proteomics data have been deposited to the ProteomeXchange Consortium via the PRoteomics IDEntifications database (PRIDE) (https://www.ebi.ac.uk/pride/archive/) partner repository with the dataset identifier PXD020950.

### Ethical Approval

For the clinical case presented here appropriate informed consent was obtained from the parents of the patient for the publication of the anonymized data. Immunohistochemistry was performed on leftover material from routine clinical procedures in the UMCU, for which informed patient consent and additional ethical approval were not required.

### Statistics

Statistical analysis was performed using GraphPad Prism 7.0 (GraphPad Software, Inc., USA). Data are presented as mean ± standard error of the mean (SEM) of at least three independent experiments performed in triplicate, unless stated otherwise. Significance was evaluated using One-way Analysis of Variance (ANOVA), or where appropriate unpaired two-tailed Student’s t-test was applied. P-values < 0.05 were considered significant.

## Author’s Contribution

A.J., M.J.J., and R.M. designed the study; A.M.vE., and M.R.L., performed human study and provided the patient’s clinical data; A.J., C.AGH.vG., E.A.Z., F.A.V., R.B., E.Z., and J.E.E. performed experiments; M.A, and C.R.B. provided input on experimental design, manuscript content and data representation; A.J., M.J.J., and R.M interpreted the data and wrote the paper with all co-authors’ assistance; M.J.J. and R.M. provided supervision.

## Acknowledgements

This work was financially supported by a grant from the Dutch Kidney Foundation (grant nr.150KG19 and KSTP12-010) and the stofwisselkracht (Druggable Targets In Lysosomal V-ATPase Dysfunction).

---

**Supplementary figure S1.**
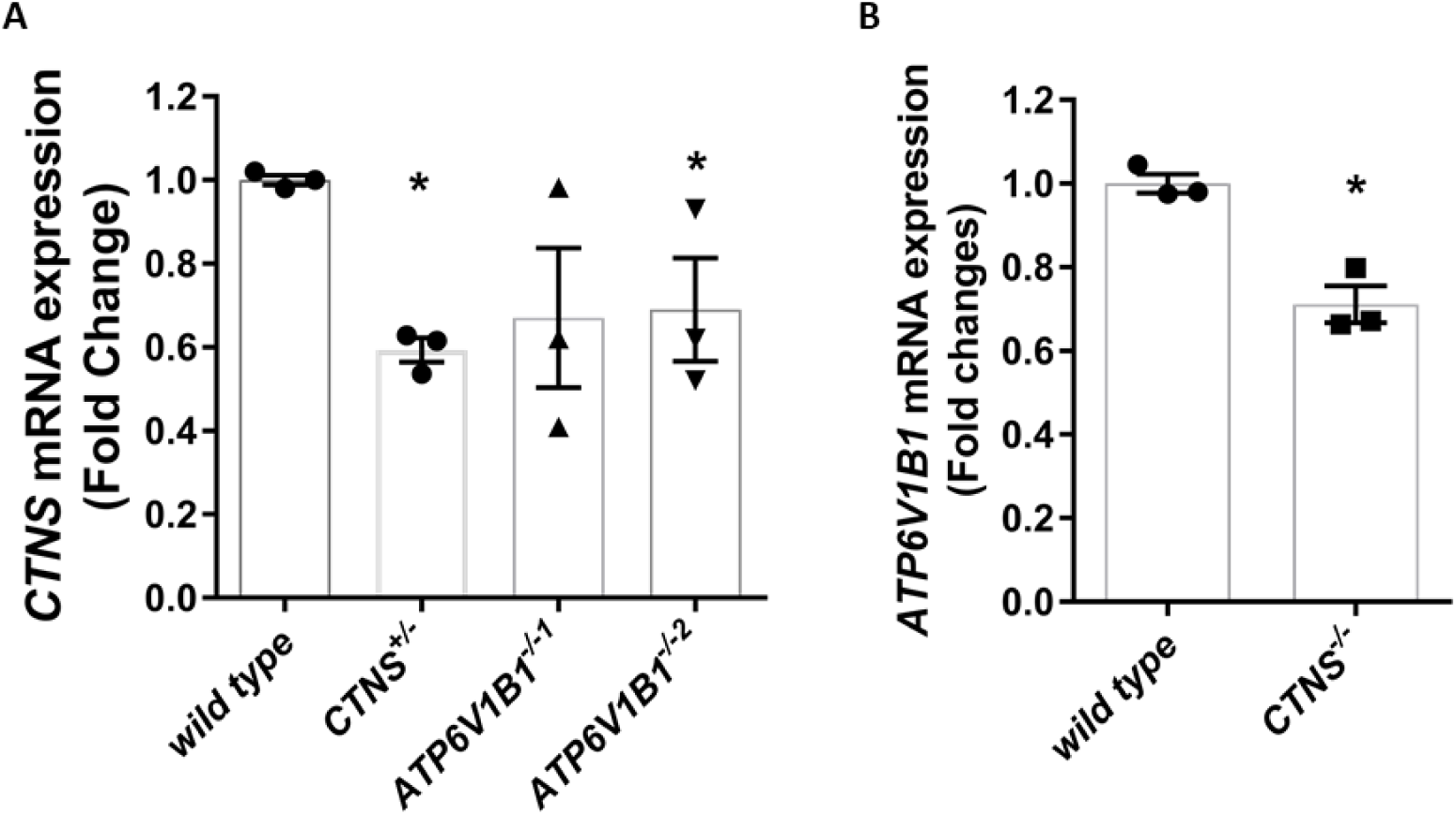
The lysosomal V-ATPase B1 interacts with cystinosin. **(A)** *CTNS* mRNA expression in two clones of *ATP6V1B1^-/-^* and *CTNS^+/-^* cells was measured using real-time quantitative polymerase chain reaction, with ribosomal protein S13 (RPS-13) as a reference gene, normalized to wild type (control) cells. **(B)** The levels of *ATP6V1B1* mRNA expression in *CTNS^-/-^* and control cells. Data are expressed as the mean ± SEM of at least three independent experiments. Statistical analysis was performed using one-way analysis of variance (ANOVA) followed by Dunnett’s multiple comparisons test. P-values < 0.05 were significant.

