## Supplementary figures and images for "The lysosomal V-ATPase B1 subunit in renal proximal tubule is linked to nephropathic cystinosis"

### Supplementary figure S1

*Supplementary figure S1*

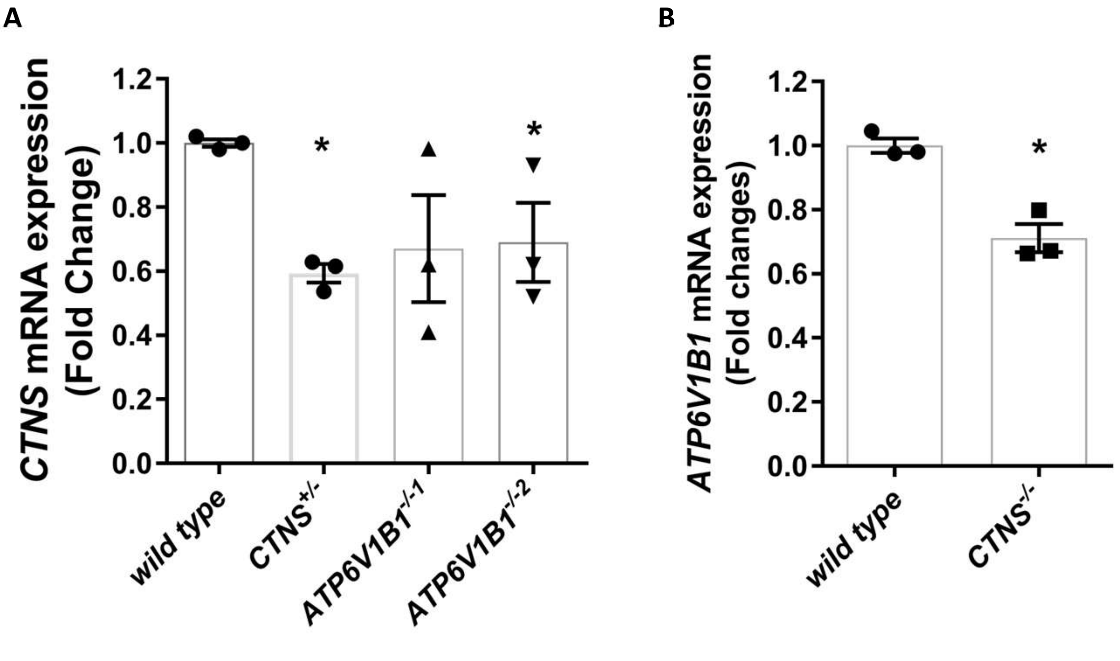
